# Evolution favors aging in populations with assortative mating and strong pathogen pressure

**DOI:** 10.1101/287664

**Authors:** Peter Lenart, Julie Bienertová-Vašků, Luděk Berec

## Abstract

Since at first sight aging seems to be omnipresent, many authors to this very day regard it as an inevitable consequence of the laws of physics. However, studies published in the past two decades have conclusively shown that a number of organisms do not age, or at least do not age on a scale comparable with other aging organisms. This disparity leads us to question why aging evolved in some organisms and not in others. We thus present a mathematical model which simulates evolution in a sexually reproducing population composed of aging and non-aging individuals. We have observed that aging individuals may outcompete non-aging individuals if they have a higher starting fertility or if the main mating pattern in the population is assortative mating. Furthermore, stronger pathogen pressure was found to help the aging phenotype when compared to the non-aging phenotype. Last but not least, the aging phenotype was found to more easily outcompete the non-aging one or to resist the dominance of the latter for a longer period of time in populations composed of dimorphic sexually reproducing individuals compared to populations of hermaphrodites. Our findings are consistent with both classical evolutionary theories of aging and with evolutionary theories of aging which assume the existence of an aging program. They can thus potentially work as a bridge between these two opposing views, suggesting that the truth in fact lies somewhere in between.

**Significance Statement:** This study presents the first mathematical model which simulates the evolution of aging in a population of sexually reproducing organisms. Our model shows that aging individuals may outcompete non-aging individuals in several scenarios known to occur in nature. Our work thus provides important insight into the question why aging has evolved in most, but not all, organisms.

## Introduction

Biological aging is defined as the age-dependent increase in the risk of death (1). And while aging is certainly a widespread process, we now know that it is far from universal (2–6). This is extremely interesting from the perspective of evolutionary biology, since it shows that evolution may produce non-aging or at least extremely slowly aging species. A proper understanding of why most species have evolved to age while some species have instead evolved to stay forever young may also help reveal how the aging process works and even how it can be modulated.

The first inquiries into the evolution of aging took place in the late 19th century (7). However, only in 1951 did Peter Medawar postulate that the force of natural selection declines with age, thus providing a key impulse which led to the formation of currently mainstream “classical” evolutionary theories of aging (8). Subsequently, Medawar’s ideas were expanded by Williams who in 1957 proposed the hypothesis of antagonistic pleiotropy, stating that evolution will select genes which are beneficial in early life even if they are deleterious later on (9). The second key expansion of Medawar’s ideas, known as the disposable soma theory, was formulated in the 1970s (10, 11). The disposable soma theory proposes that strong evolutionary pressure is necessary to maintain the integrity of germ cells but comparatively very weak pressure is essential for maintaining somatic tissues. As a result, somatic tissues are not maintained well enough, which results in time-dependent deterioration, i.e. aging (12).

Classical evolutionary theories of aging consider aging to be an inevitable byproduct of evolution, and, while there are logically consistent, they are unable to explain the existence of non-aging species. For this and other reasons, several authors have proposed that aging is directed by an aging program (13–20). However, since the classical view of the evolution of aging precludes the evolution of such a program, alternative views of how aging might be favored by natural selection have also been suggested (15, 18, 21). For example, Josh Mitteldorf proposed that aging has evolved in order to stabilize population dynamics (14, 21, 22). Furthermore, several mathematical models have demonstrated that, under specific conditions, aging can be selected by evolution (23–26). However, these models have limited applicability. Some assume that even without aging fertility decreases in an age-dependent manner (23) or that competitive fitness declines with age (25) and thus in a certain sense require aging for the evolution of aging. More importantly, the methodology, assumptions, and conclusions of each of these models have been called into question and challenged in some detail (27). Furthermore, though the results of these models are interesting, they all simulate the evolution of aging in asexually reproducing organisms; their results thus have limited application for the evolution of aging in sexually reproducing organisms.

We have previously proposed a theoretical verbal model (28) which states that although aging is, in essence, inevitable and results from damage accumulation rather than from a specific program, the actual rate of aging in nature may still be adaptive to some extent. Our verbal model also briefly describes how aging individuals might outcompete seemingly non-aging individuals under strong pathogen pressure or rapidly changing environmental conditions. In this article, we derive a complex mathematical model which simulates competition between aging and non-aging individuals in a sexually reproducing population formed either by hermaphrodites or by true sexuals (males and females) under defined pathogen pressure.

## Methods

We begin with a model population composed of *N* hermaphrodites of two phenotypes: aging and non-aging. We develop an agent-based simulation model that allows phenotypes of all individuals to be modeled explicitly and their competitive dynamics to be followed over time. Time is discrete, with the time step corresponding to the age increment of 1 and with all relevant rates and probabilities defined on a per time step basis. Simulations are run for *T* time steps and each model scenario (see below) is replicated 20 times. All model parameters and variables are summarized in Table 1. The model is subsequently extended to true sexually reproducing populations composed of males and females. It should be noted that this model is a modification of the classical models of the evolution of sexual reproduction (29–32) which also takes into account individual age and aging and non-aging phenotypes.

**Table 1:**
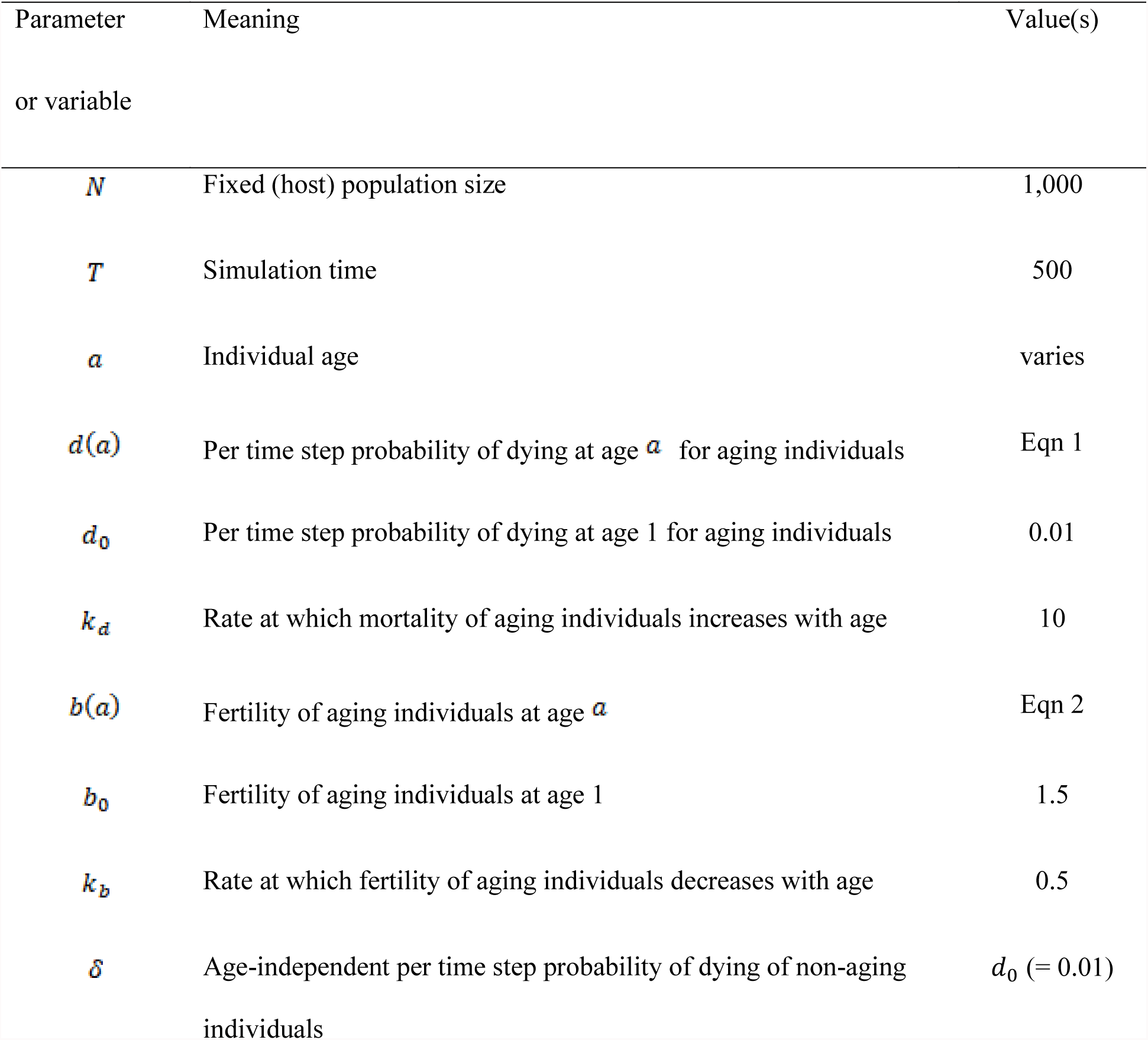
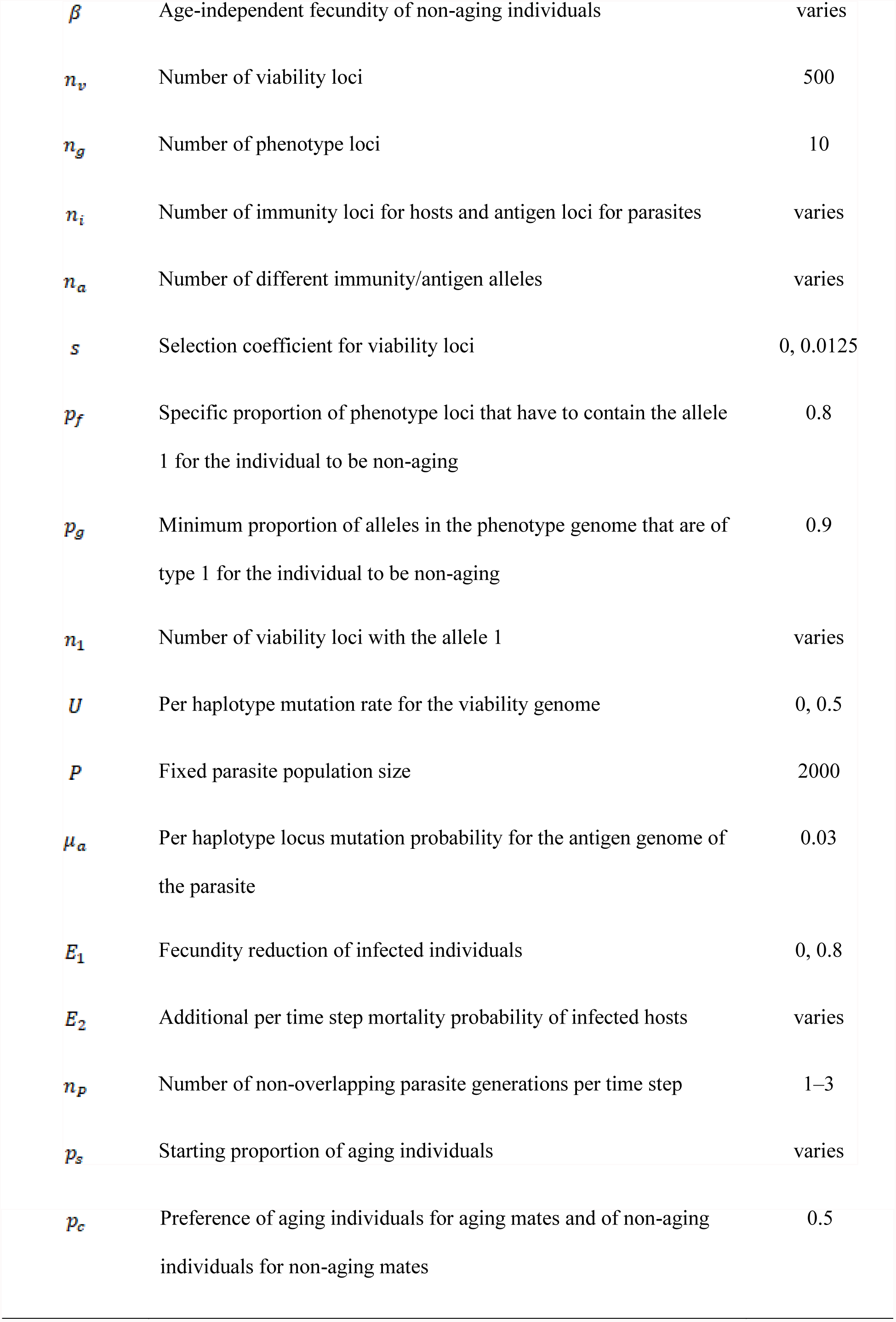
Default parameter and variable settings used in the hermaphrodite model.

### Demography

Aging and non-aging phenotypes differ according to demographic rates. Each individual is characterized by age *a*. The probability that an aging individual dies during a time step increases with age *a* as

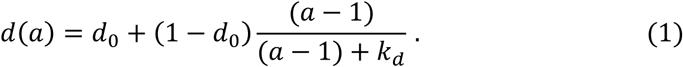

Similarly, the fecundity of an aging individual decreases with age *a* as

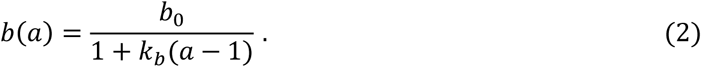

The (*a −* 1) term in equations (1) and (2) is introduced so that mortality and fecundity equal *d*_0_ and *b*_0_ for individuals of age *a* = 1. Newborn mortality in the first year of life is included in parameter *b*_0_ and equations (1) and (2) are thus applicable to individuals of age *a* > 0. A non-aging individual dies during a time step with an age-independent probability *δ* and produces *β* offspring in each time step, irrespective of age.

### Individuals

In addition to age, each individual is characterized by three genome portions. The viability genome, composed of *n*_*v*_ loci, serves to quantify the impact of selection and resulting offspring fitness (see below). The phenotype genome, composed of *n*_*g*_ loci, serves to determine whether a newborn individual belongs to the aging or non-aging phenotype (see below). Finally, the role of the immunity genome, composed of *n*_*i*_ loci, is to check for an individual’s resistance to a parasitic infection (see below). All three genome portions are assumed haploid. When we consider true sexuals later on, the viability and phenotype genomes are assumed diploid. Also, individuals are characterized as susceptible or infected (all are initially susceptible, and all are born susceptible).

### Genetic structure

Any viability genome locus may feature one of two allele types, with 0 and 1 denoting neutral and deleterious alleles respectively. In the case of a viability genome, the fitness of an individual is calculated as

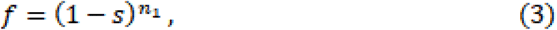

where *s* is the selection coefficient and *n*_*1*_ is the number of loci with the deleterious allele 1.

Any phenotype genome locus may also feature one of two allele types, with 0 and 1 denoting alleles contributing to the aging and non-aging phenotype respectively. While there are many viable options how to determine the aging and non-aging phenotype from a particular phenotype genome, we assume that a number of loci must be in harmony in order to produce a non-aging phenotype; as a result, we have opted for adopting a threshold model. In particular, two conditions must be satisfied for an individual to be considered non-aging: (1) the specific proportion py of phenotype loci must contain the allele 1, and (2) the proportion of *p*_*f*_ all phenotype genome type 1 alleles must exceed a given threshold value *p*_*g*_ > *p*_*f*_.

Finally, in the immunity genome, alleles denoted as *0*,…*n*_*a*_ − 1 represent *n*_*a*_ alternative variants of the immunity allele. The viability, phenotype and immunity genomes of each offspring are determined following the free recombination of their parents’ genomes and a random choice of one of two resulting genomes.

The per haplotype locus probability of a deleterious mutation for the viability genome is *U*/*n*_*v*_ (i.e. the actual number of deleterious mutations per large enough haplotype is a Poisson variable with mean *U*). No phenotype and immunity genome mutations are assumed to occur.

For the viability genome, individuals are initialized with *U/s* deleterious alleles. The population is thus initialized at the mutation-selection equilibrium for the number of deleterious mutations. For the phenotype genome, alleles at all loci of aging individuals are initially of type 0 and alleles at all loci of non-aging individuals are initially of type 1. Initially, an allele is randomly selected for each locus in the immunity genome.

### Parasites

We simulate a parasite population of a fixed size *P.* Individual parasites are characterized by the haploid antigen genome, with *n*_*a*_ alleles denoted as 0,…,*n*_*a*_ − 1 that are initially randomly distributed across the loci. Genetic variation at the antigen loci in the parasite population is maintained by setting the per locus mutation rate to *μ*_*a*_ per parasite generation. When a mutation at a locus occurs, the existing allele is replaced by an allele randomly chosen from the *n*_*a*_ alleles 0, …,*n*_*a*_ − 1. Parasites reduce the fecundity of the infected individuals by a certain factor *E*_*1*_. Moreover, the infected individuals die during the time step with an extra probability *E*_*2*_. Since parasite life cycles are commonly much faster than those of their hosts, we assume there are *n*_*p*_ non-overlapping parasite generations per time step.

### Time step

Within any time step, parasitism is applied first to all host individuals. During each parasite generation, hosts are drawn sequentially in a random order and each exposed to a randomly selected parasite. We use a matching alleles model of infection genetics (29–34) to establish whether infection actually occurs in the host: if the immunity genome of the host and the antigen genome of the parasite match exactly at all loci, the parasite establishes infection in the host and is placed into a pool of successful parasites, otherwise it dies. Hosts in which infection is established are marked as infected. At the end of each parasite generation, *P* new parasites appear in the environment by drawing parasite individuals at random (with replacements) from the pool of successful parasites. Following the introduction of a new parasite, its antigen alleles are allowed to mutate to any of the *n*_*a*_ alleles 0, …,*n*_*a*_ − 1, each with a certain probability *μ*_*a*_. To examine the impact of parasitism, selected simulations are run without this parasitic phase.

Upon calculating infection prevalence among aging and non-aging individuals, the parasitic phase is followed by host demography. First, individuals are allowed to mate and reproduce. There is a preference *p*_*c*_ of aging individuals for aging mates and of non-aging individuals for non-aging mates, but otherwise mates are chosen randomly. We note that *p*_*c*_ = 0.5 indicates no preference and thus a random choice of mates by all host individuals regardless of phenotype. Upon mating, a Poisson-distributed number of offspring are produced, with mean *b(a)* and *β* for aging and non-aging individuals, respectively, reduced by a factor 1 − *E*_1_ if hosts are infected. The offspring are born susceptible, with age 1 (as we emphasized earlier, fecundity already accounts for first-year mortality), and the viability and phenotype genomes of each offspring are determined (see above). The probability that an offspring survives the embryonic phase is given by its fitness *f* (which equals 1 unless the selection coefficient *s* is positive). The phenotype (aging or non-aging) of each surviving offspring is then determined using the above-described threshold model.

Background mortality of other than newborn hosts then occurs, followed by the additional mortality probability *E*_*2*_ of infected hosts. The age of all individuals is augmented by 1 and we record the numbers of aging and non-aging individuals and the previously calculated infection prevalence among aging and non-aging individuals. Eventually, a maximum of *N* individuals is randomly selected to form the population appearing at the beginning of the next time step.

### Extended model version with sexual dimorphism

In order to adapt our model for true sexuals with explicit males and females, some of its components were replaced and new elements were introduced. Aside from these modifications, the principles of the extended model mirror those of the haploid hermaphrodite model. In order to adapt the model, we first replace hermaphrodites with females and males. Among other things, this means that females select their mating partners from among males and that only females produce offspring. Since hermaphrodites need to allocate some amount of resources to both male and female functions, then – relative to females – their fertility is assumed to be reduced, as much as by one half (35). Therefore, to fairly compare simulations of hermaphrodites and true sexuals, we assume female fertility to be twice that of hermaphrodites. On the other hand, the model assumes that both females and males have identical mortality patterns, depending on whether they are aging or non-aging.

The viability and phenotype genomes in the sexually dimorphic population are diploid. In the viability genome, deleterious alleles are assumed to be relatively recessive. That is, if selection occurs, the fitness of an individual is calculated as

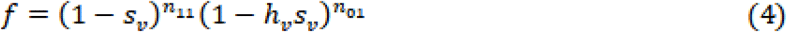

where *s*_*v*_ and *h*_*v*_ are selection and dominance coefficients, respectively, and *n*_*11*_ and *n*_01_ are the numbers of loci homogeneous for an allele 1 and heterogeneous loci, respectively, in the viability genome. Moreover, true sexuals are initialized with *U/(hs)* deleterious mutations when *h* > 0 or with *U/s* deleterious mutations when *h =* 0. In this manner, the population is again initialized at the mutation-selection equilibrium for the number of deleterious mutations.

In the phenotype genome, non-aging alleles are assumed to be relatively recessive. Again, we adopt a threshold model where two conditions must be satisfied for the individual to be non-aging: (1) the proportion *p*_*f*_ of fixed genotype loci must be homozygous for allele 1, and (2) the proportion of alleles 1 in the entire phenotype genome must exceed a given threshold value *p*_*g*_> *P*_*f*_.

## Results

### Demographic parameters

Our first objective was to find an optimal setting for variables defining initial fertility and the proportion of aging and non-aging individuals in a population. We tested 12 different settings, as summarized in Table 2, with a simulation time of *T* = 300. In this initial simulation set, parasite infection reduced host fertility by 50% but never killed its host.

**Table 2:**
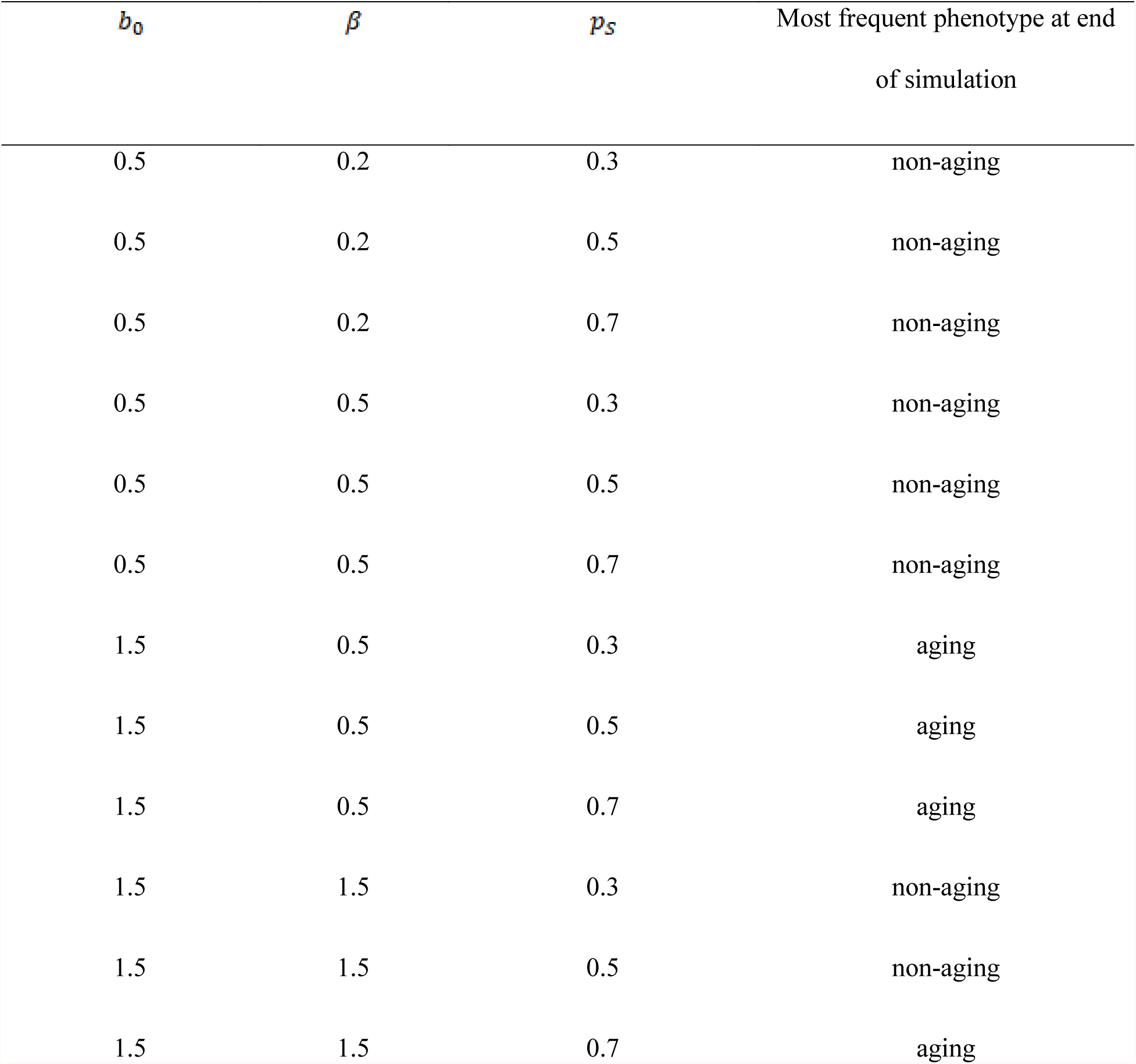
Tested settings of basic demographic parameters in our preliminary test.

The graphical representations of these simulations can be found in Fig S1. The results of these simulations suggest that aging individuals may win the competition if their initial fertility is sufficiently high relative to that of non-aging individuals. Moreover, in case the non-aging phenotype eventually prevails, aging individuals remain in the population the longer the higher their initial frequency in the population. It is also interesting to note that with an equal initial fertility of 1.5 for both aging and non-aging individuals, the aging individuals fare better than when their initial fertility of 0.5 is higher than that of non-aging individuals set at 0.2. As a consequence, both relative and absolute values of fertility appear to drive the outcome of the competition.

Next, we tested what smallest possible difference in initial fertility between aging and non-aging individuals still results in the fixation of the aging phenotype after *T* = 500 generations and with the initial proportion of aging individuals set at *p*_*s*_ = 0.5 (Fig. S1). First, we assumed no infection, set initial fertility *b*_*0*_ of aging individuals at 1.5 and tested scenarios with various lower fertilities *β* of non-aging individuals. Simulations with fertility of non-aging individuals set at 1.1 or lower always resulted in 100 % dominance of the aging phenotype after *T =* 500 generations. Furthermore, simulations with fertility of non-aging individuals set at 1.2 still, in a great majority of cases, resulted in the domination of the aging phenotype. However, higher fertility levels of non-aging individuals resulted in their domination. Thus, the minimal difference in initial fertility which enables the dominance of the aging phenotype in this specific setting is 20% of the initial fertility of the aging phenotype. However, when simulating the same scenarios with infection (*E*_*1*_ = 0, *E*_*2*_ = 0.4, *n*_*i*_ = 1, *n*_*a*_ = 8) the minimal difference in initial fertility required for the domination of the aging phenotype decreased. With infection present, the aging phenotype dominated most of the time even with the fertility of the non-aging phenotype set at 1.3.

We subsequently selected *b*_*0*_ = β = 1.5 and *p*_*s*_ = 0.5. We applied the same settings to aging and non-aging phenotypes in order to facilitate their fair comparison in different conditions. Moreover, preliminary tests indicated that initial fertilities lower than 1 compromise the ability of both aging and non-aging phenotypes to survive after factoring in pathogen pressure and selection.

### Effect of selection and mating preferences

In the next set of simulations, we tested the effect of selection and mating preferences for either the same or opposite aging phenotype. To better understand the effect of these changes, the simulations were run without factoring in infection. In contrast to our expectations, the simulation results suggest that selection is beneficial for the non-aging phenotype (Fig.1). While the figure presents only one simulation scenario, the results of the remaining scenarios were analogous.

**Figure 1:**
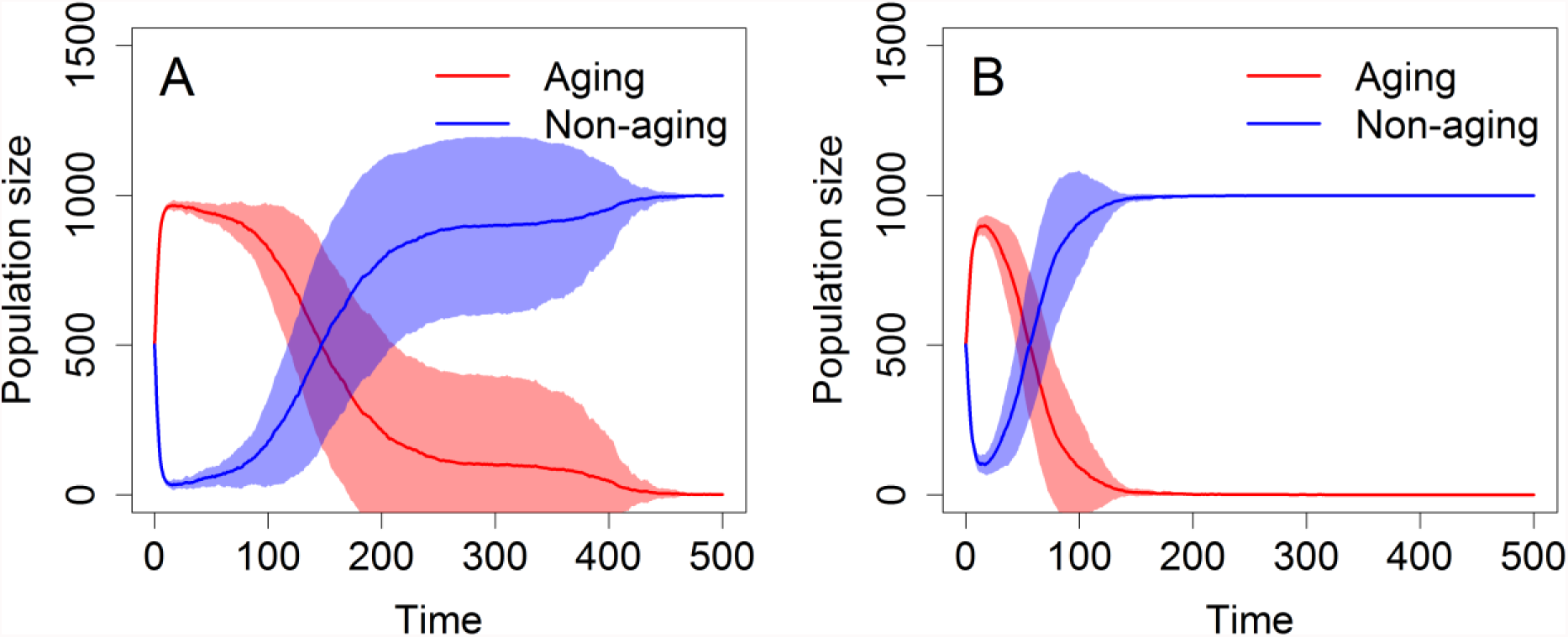
Changes in the representation of aging and non-aging phenotypes in the population under different simulation settings: A) No selection, random mating; B) Selection and random mating.

As a result of our threshold approach to modeling the inheritance of aging, we observed a massive increase in the number of aging individuals in the first few generations in every simulation (Fig.1 and additional figures below). This is caused by the fact that at the start of every simulation, the phenotype loci of all aging and non-aging individuals contained solely aging or non-aging alleles, respectively.

Similarly, the results of simulations with biased mating preferences have also exceeded our expectations. Mating preferences were tested with a wide range of *P*_*c*_ values. To elucidate their meaning, e.g. *p*_*c*_ = 0.3 corresponds to a 70% probability of mating with the opposing phenotype (aging with non-aging and vice versa; disassortative mating) while *p*_*c*_ = 0.7 corresponds to a 70% probability of mating with the same phenotype (assortative mating). In general, a preference for the opposing phenotype (*p*_*c*_ < 0.5) leads to the formation of a coexistence equilibrium in the proportion of aging and non-aging phenotypes (Fig.2). On the other hand, while *p*_*c*_ > 0.5 but smaller than 0.9 favors the aging phenotype, *p*_*c*_ = 0.9 (and also higher values) results in fast and stable dominance of the non-aging phenotype (Fig.2). Thus, unless the preference for the same phenotype is not very high, we observe the rapid dominance of the aging phenotype.

**Figure 2:**
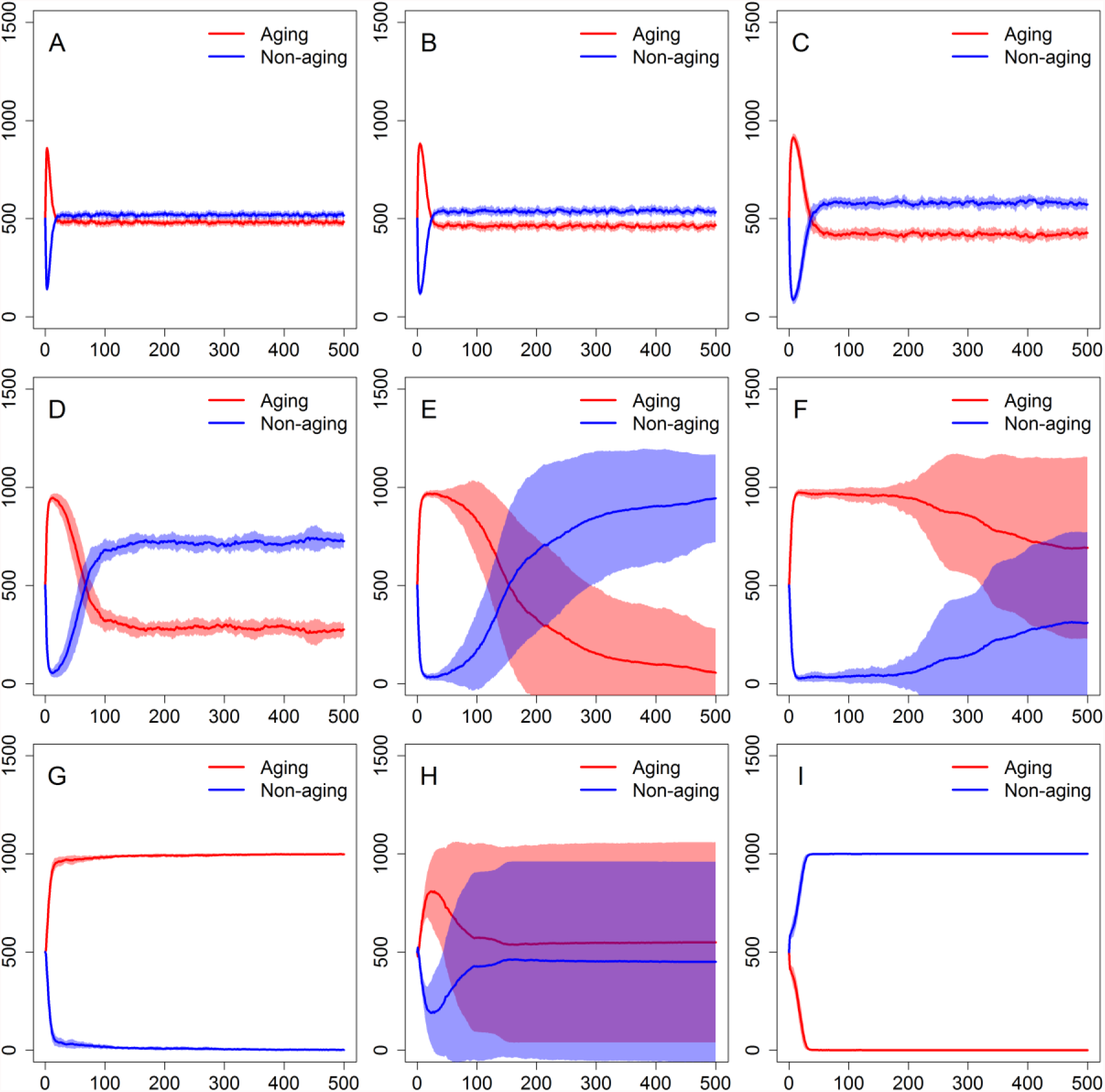
Changes in the representation of aging and non-aging phenotypes in the population under different simulation settings: A) *P_c_* = 0.1; B) *P_c_* = 0.2; C) *P_c_* = 0.3; D) *P*_*c*_ = 0.4; E) *P*_*c*_ = 0.5; F) *p*_*c*_ = 0.6; G) *P*_*c*_ = 0.7; H) *P*_*c*_ = 0.8; I) *P*_*c*_ = 0.9.

### Pathogen settings

Next, we tested the effect of different infection settings: a decline in fertility (*E*_1_), an increase in mortality (*E*_*2*_), and the number of immunity/antigen loci (*n*_*i*_,) and alleles (*n*_*a*_).

Interestingly, relative to the no-infection case (Fig.2E), the infection-caused decline in fertility accelerates the domination of the non-aging phenotype while an increase in mortality has the opposite effect (Fig.3). Surprisingly, the higher number of immunity/antigen alleles seems to slightly accelerate the domination of the non-aging phenotype (results not shown). Thus, in a subsequent step, we tested the effect of the increase in the number of immunity/antigen loci. However, increasing the number of immunity/antigen loci without changing the number of immunity/antigen alleles seems to have only a marginal effect on the domination of the non-aging phenotype (results not shown).

**Figure 3:**
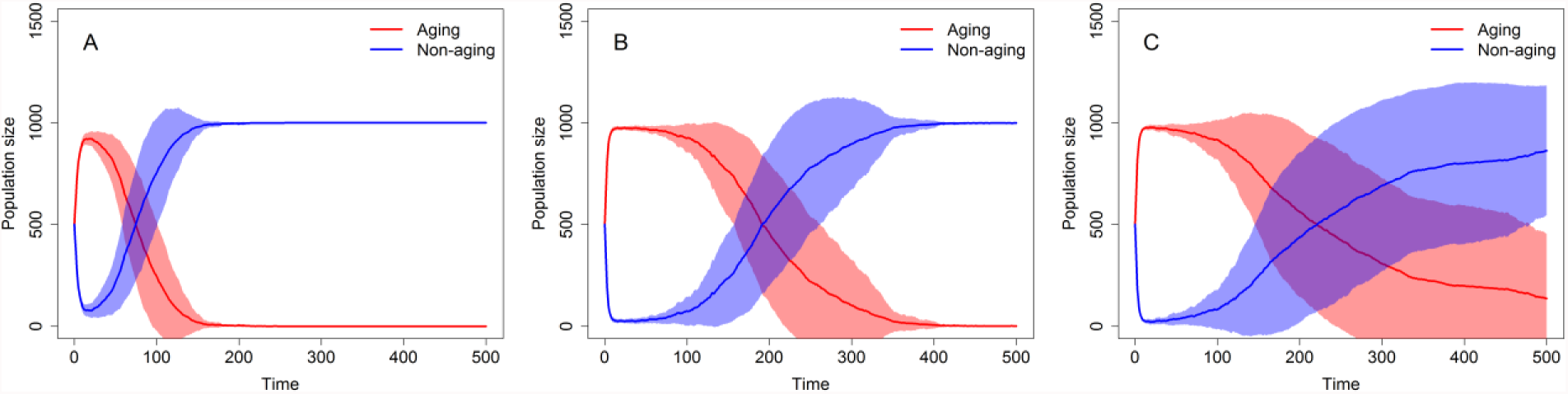
Changes in the representation of aging and non-aging phenotypes in the population under different simulation settings: A) *n*_*a*_ =4, *E_1_* = 0.8 and *E*_*2*_ = 0; B) *n*_*a*_ = 4, *E*_*1*_ = 0 and *E*_*2*_ = 0.2; C) *n_a_=* 4, *E*_*1*_ = 0 and *E*_*2*_ = 0.4.

We also tested the effect of pathogen population size (*P*) and the number of pathogen generations per time step (*n*_*P*_). Tested pathogen population sizes included *P =* 1,000 and *P =* 3,000 in contrast with the default setting of *P =* 2,000. One to three pathogen generations per time step were simulated. All scenarios were tested with both 20% and 40% infection-related mortality settings.

Greater pathogen population size delayed the dominance of the non-aging phenotype (Fig.S2). The number of pathogen generations had a similar effect; every additional pathogen generation was found to postpone the dominance of the non-aging phenotype (Fig.4).

**Figure 4:**
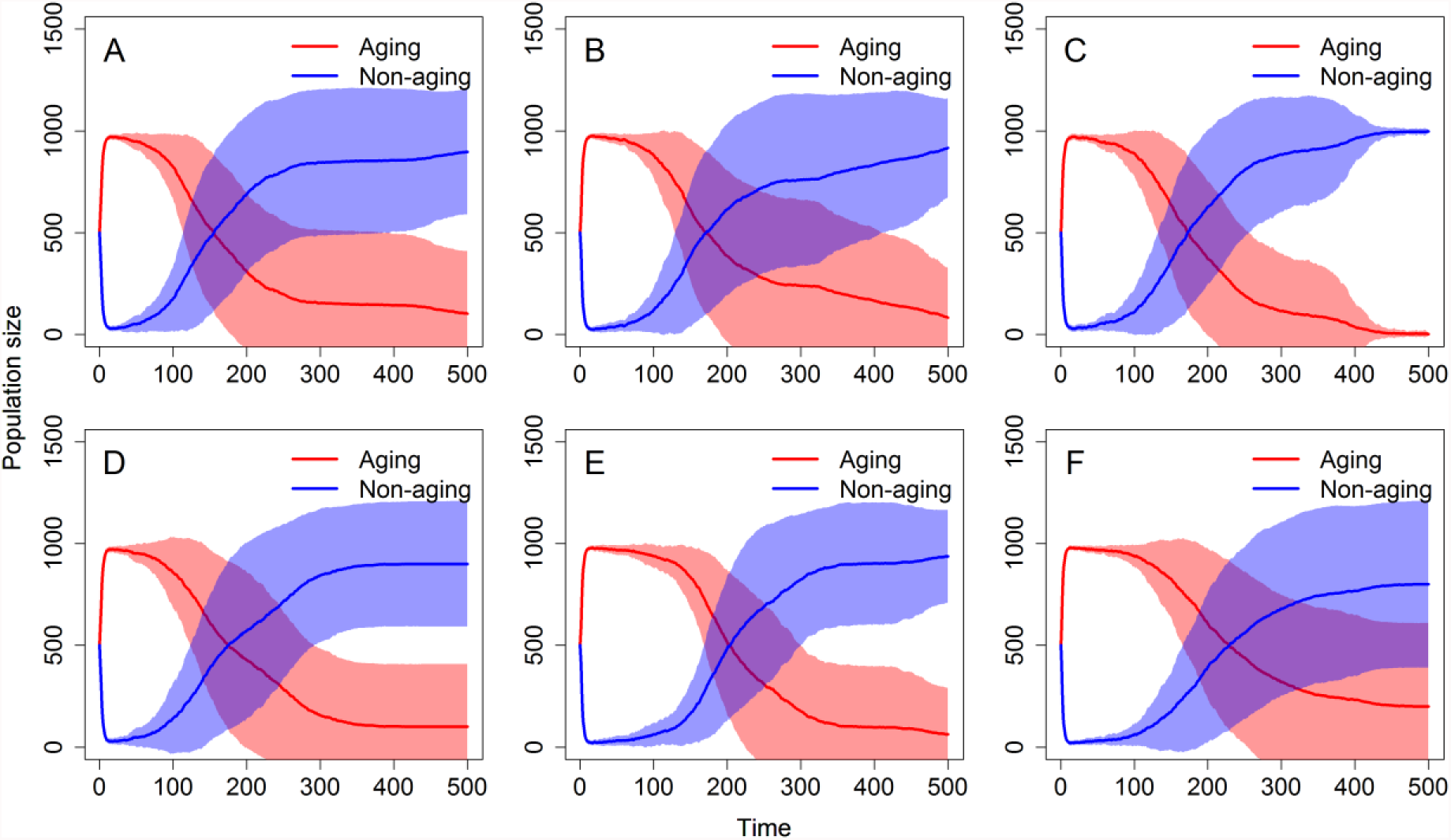
Changes in the representation of aging and non-aging phenotypes in the population under different simulation settings. A) *E_2_* = 0.2, *n_p_* = 1; B) *E_2_* = 0.2, *n_p_* = 2; C) *E_2_* = 0.2, *n_p_* = 3; D) *E*_*2*_ = 0.4, *n*_*p*_ = 1; E) *E*_*2*_ = 0.4, *n*_*p*_ = 2; F) *E*_*2*_ = 0.4, *n*_*p*_ = 3.

### Multi-factorial scenarios

In nature, nothing works in isolation. Thus, as the final test of our simplified hermaphrodite population model, we selected several multi-factorial scenarios combining several of the previously tested situations. The tested scenarios are summarized in Table 3.

**Table 3:**
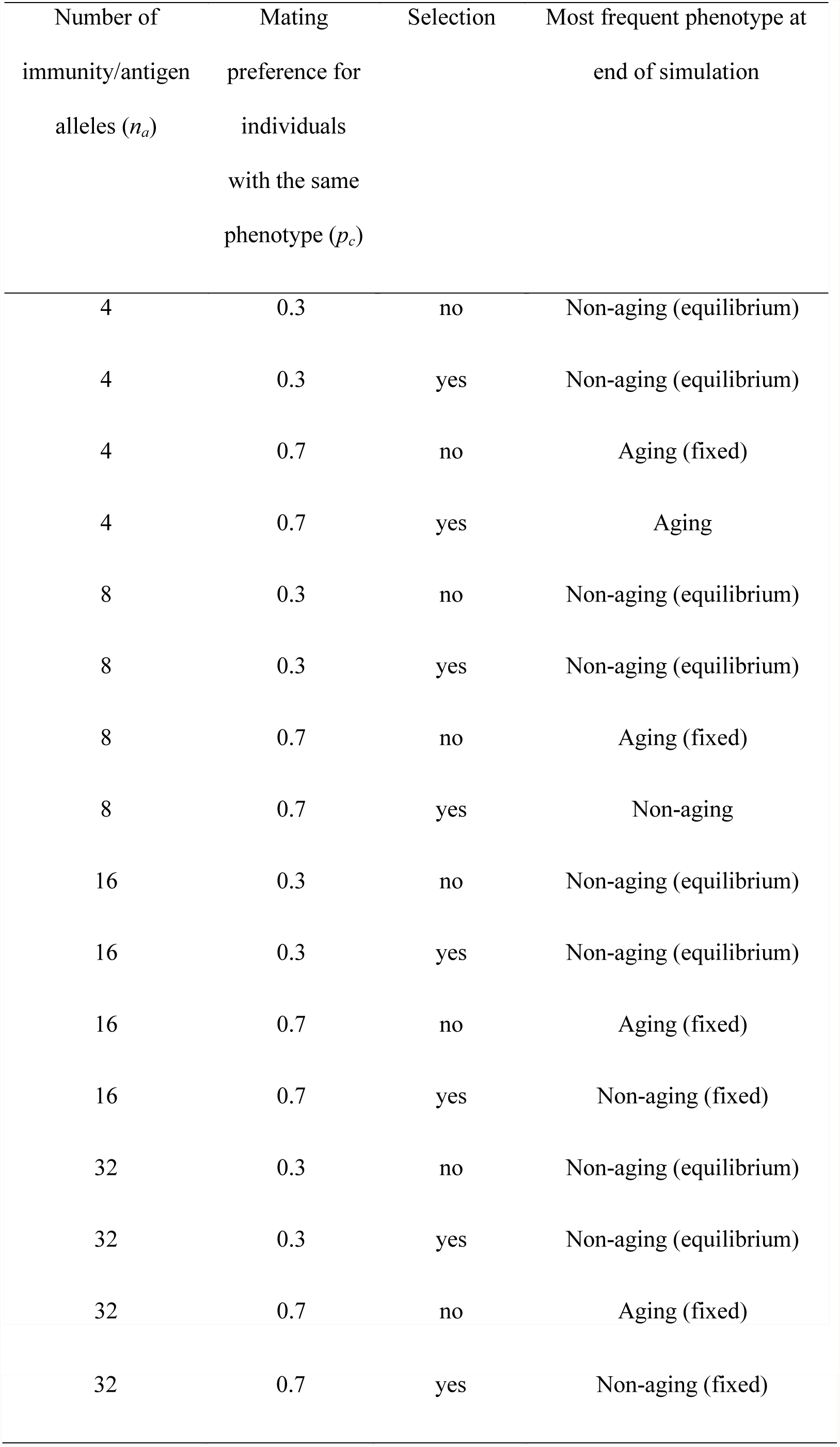
Multi-factorial scenarios for the hermaphrodite population. The phenotype at the end of the simulation was considered fixed if it accounted for over 95% of the population for at least the last 100 generations. If no phenotype was fixed or in equilibrium at the end of the simulation, we noted the more frequent phenotype. The phenotypes at the end of the simulation were in equilibrium if they cycled around same values for at least 300 generations. In case selection was present, a value of *s* = 0.0125 was used.

These results show that the aging phenotype can be fixed in populations with assortative mating and that this effect can be disrupted by selection. On the other hand, the number of immunity/antigen alleles does not appear to have any effect. These results thus confirm the predictions of previous simulations focusing on one variable per system at a time.

### Sexual dimorphism

To test whether the results of our simulations are applicable to sexually dimorphic organisms, we developed an extended version of our model which includes diploid males and females instead of haploid hermaphrodites. First, we tested the effect of different initial fertility values between aging and non-aging individuals (Fig.5). Overall, the results of the sexual model were in agreement with previous simulations; even a relatively small difference in initial fertility in favor of the aging phenotype was observed to lead to its long-term domination. However, in contrast with previous simulations, the time required for the non-aging phenotype to dominate was extremely long even when no difference in initial fertility was set. Even after 1,000 generations the aging phenotype was more frequent in most simulations. A dimorphic sexual population is thus vastly more favorable for the aging phenotype then a population composed of hermaphrodites.

**Figure 5:**
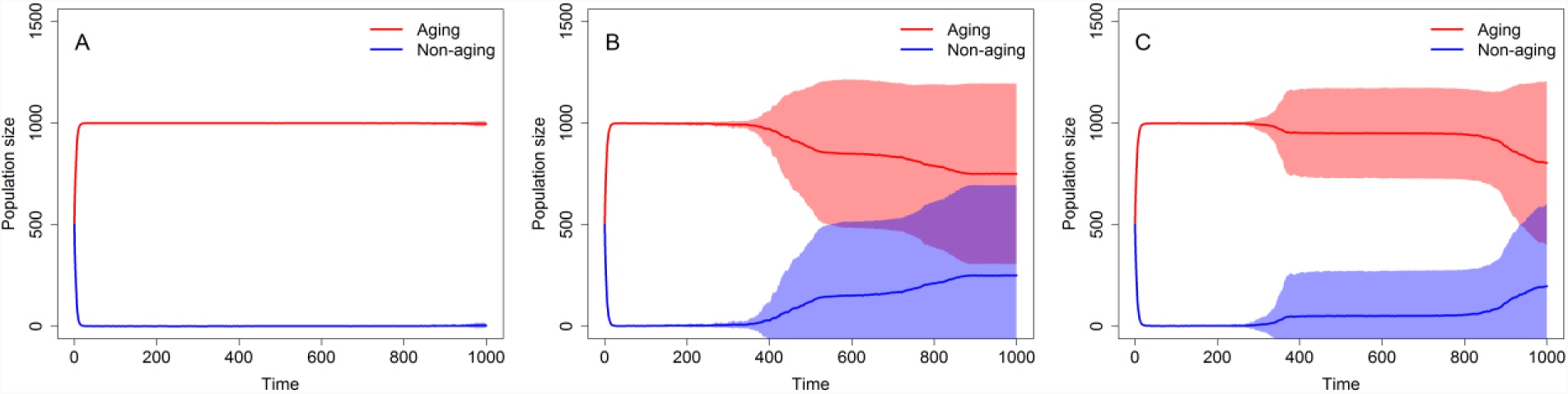
Changes in the representation of aging and non-aging phenotypes in the population with true sexuals under different simulation settings: A) *β* = 2 × 1.3; B)*β* = 2 × 1.4; C). Initial aging phenotype fertility is *b*_0_ = 2 × 1.5.

In the second series of scenarios, we tested the effect of pathogen pressure on the extended model (Fig.S3F-J). In contrast with previous simulations, the infection slightly reduced the domination of the aging phenotype. This effect was most notable in simulations with a higher number of antigen/immunity alleles and loci. We hypothesize that incompatibility with the results of simulations in the basic version of our model might be caused by the increase in initial fertility introduced to the extended version of the model in order to balance the reduction of a number of offspring-producing individuals caused by introducing males. We tested this hypothesis by testing the effect of infection with the same initial fertility as in previous simulations (Fig.S3A-E). After this adjustment, the results mirrored those derived from the hermaphrodite model version. Most notably, pathogen pressure provided a clear benefit for the aging phenotype and delayed the dominance of the non-aging phenotype. Furthermore, the benefit of pathogen pressure for the aging phenotype was most notable with a lower number of immunity/antigen alleles and loci decreasing rapidly with their increase; this effect of immunity/antigen alleles and loci was much more pronounced than in the basic version of our model. We also compared results at both initial fertility settings (Fig.S3) with the same settings in the hermaphrodite population (Fig.S4). The comparison showed that while higher initial fertility is beneficial for the aging phenotype in both versions of the model, the extended version of the model is more favorable for the aging phenotype regardless of the initial fertility setting.

We recall that doubling female fertility relative to that of hermaphrodites corresponds to the assumption that hermaphrodites allocate the same amount of resources to male and female functions, whereas the equal fertilities of females and hermaphrodites reflect an extreme situation where the male function of hermaphrodites bears no costs. Thus, in order to find out how the population dynamics differ for hermaphroditism and sexual dimorphism when the cost of males is low, we compared simulations with b_0_ = *2 ×* l.5 hermaphroditic (Fig.S4 F-J) and fully sexual (Fig.S3 F-J) models. However, to assess the difference when the cost of males is high, we compared simulations with b_0_ = 1.5 for hermaphrodites (Fig.S4 A-E) with simulations b_0_ = 2 × 1.5. for the fully sexual model (Fig.S3 F-J). These comparisons suggest that as the cost of the male function in hermaphrodites increases, so does the positive effect of parasites on the persistence of the aging phenotype.

In the third series of scenarios, we tested the effect of mating preferences (Fig.6). In this case the results were in good agreement with the results of the hermaphridite model version. The *p*_*c*_.<0.5. led to the formation of a coexistence equilibrium in the frequency of the aging and non-aging phenotypes, while *p*_*c*_ > 0.5 but smaller than 0.9 lead to permanent domination of the aging phenotype. Finally, *p*_*c*_ = 0.9 resulted in the fast and stable dominance of the non-aging phenotype. Nevertheless, even though the results of these series of simulations showed the same trend established previously in the hermaphrodite version of our model, it was still evident that the sexually dimorphic population is much more favorable in the case of the aging phenotype.

**Figure 6:**
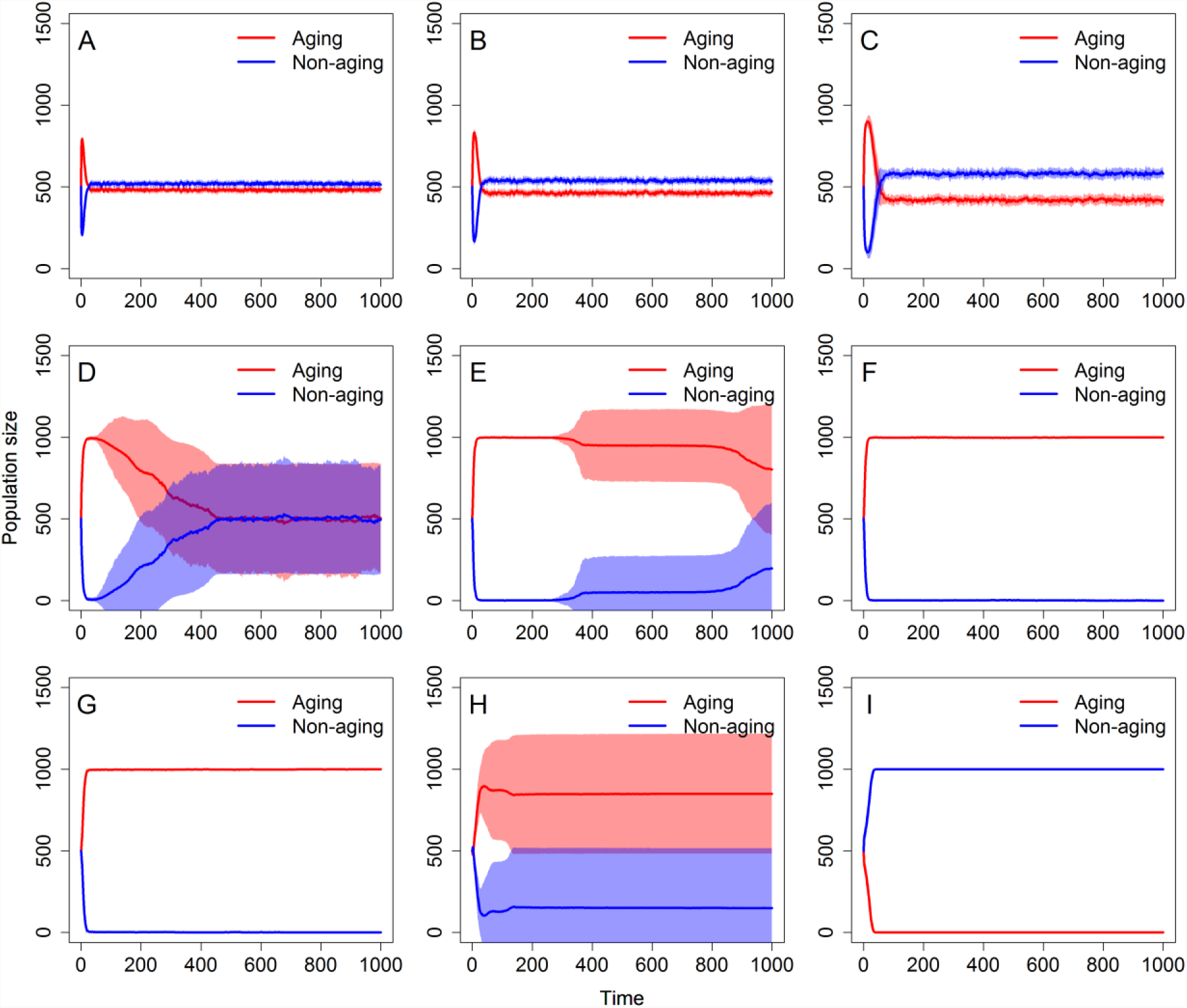
Changes in the representation of aging and non-aging phenotypes in the population of true sexual under different simulation settings: A) *P*_*c*_ = 0.1; B) *P_c_* = 0.2; C) *P_c_* = 0.3; D) *P*_*c*_ = 0.4; E) *P*_*c*_ = 0.5; F) *P*_*c*_ = 0.6; G) *P*_*c*_ = 0.7; H) *P*_*c*_ = 0.8; I) *P*_*c*_ = 0.9.

## Discussion

To the best of our knowledge, this study is the first to present a mathematical model of the evolution of aging in sexually reproducing organisms. Our results offer several insights into the evolution of aging, and while all of them are potentially interesting, four deserve special attention. First, our simulations show that higher initial fertility can outweigh a decrease in fitness associated with aging. Second, coevolution with a pathogen, particularly one that is deadly and has a quick reproduction cycle, can profoundly extend the time required for a non-aging phenotype to outcompete aging individuals. Our third – and arguably most interesting – result indicates that assortative mating (with regard to the aging phenotype) by itself leads to quick and permanent domination of the aging phenotype, while disassortative mating leads to the formation of an equilibrium in the number of aging and non-aging individuals. Fourth, the aging phenotype fares much better in a model with sexual dimorphism than in a model with sexually reproducing hermaphrodites.

Our first core finding is that the higher initial fertility of aging individuals may lead to the fixation of the aging phenotype. The main strength of this finding is that it can be easily integrated into classical theories of aging. Since both the theory of antagonistic pleiotropy (9) and the disposable soma theory (11) predict that lifespan and reproduction are in a trade-off relationship, it makes sense to assume that aging individuals would reproduce much faster than non-aging ones. This result alone is therefore capable of explaining why aging is so widespread in nature. Non-aging may thus simply evolve in organisms where it does not significantly reduce fertility or, in other words, where the fecundity cost to non-aging is low.

On the other hand, our second core finding, which states that strong pathogen pressure slows the pace at which non-aging individuals outcompete aging ones, is in accordance with some alternative views of the evolution of aging. For example, in our previous work, we have invoked the Red Queen hypothesis and proposed that a faster rate of aging may be selected because aging could enable faster adaptation to changing conditions, most notably coevolution with a pathogen (28). Furthermore, others have also invoked the Red Queen hypothesis in a similar manner with regard to the evolution of aging (24). Our results thus support the notion that the presence of pathogens may be one of the forces driving the evolution of aging. However, our simulations modeling competition between aging and non-aging individuals show that pathogen pressure is not strong enough to facilitate a fixation of the aging phenotype. Nevertheless, competition between aging and non-aging individuals is a model situation which is in essence extreme and it is thus possible that pathogen pressure may have a more profound effect on competition between individuals with different rates of aging. Furthermore, even in our extreme scenario, pathogen pressure can lead to the fixation of the aging phenotype if combined with other effects.

Our third core finding – which is also arguably the most surprising result of our simulations – is that assortative mating, if not strong enough, leads to the fast and permanent fixation of the aging phenotype. In essence, assortative mating constitutes a barrier in gene flow and as such may be the result of causes other than an active preference for the opposing phenotype. For example, if we consider a large population composed of smaller spatially divided subpopulations which are predominantly but not exclusively composed of individuals with the same phenotype, then even if all individuals chose their mating partners randomly, the population as a whole will most probably mate assortatively because individuals from different subpopulations meet only occasionally. In such a case, assortative mating is the outcome of geography. In other words, this effect may be caused by many different situations, and it is thus likely to play some role in the evolution of aging across the tree of life. However, our results also suggest that very strong assortative mating (90% preference for the same phenotype) will, unlike less strict assortative mating, favor the non-aging instead of the aging phenotype. While this 180-degree turn in the effect of assortative mating may seem surprising, it has a rather simple explanation. In particular, very strong assortative mating will generate two almost separate populations, which leads to the rapidly achieved dominance of the non-aging phenotype since it on average produces more offspring per generation than the aging one. Nevertheless, although natural populations with very strong assortative mating where almost 100% of mating occurs between similar phenotypes do exist (36), assortative mating is not nearly as strong in most species and for most phenotypes. Furthermore, the benefit of very strong assortative mating for the non-aging phenotype depends on equal initial fertility of the aging and non-aging phenotype. However, it is highly likely that the aging phenotype would have a higher initial fertility. In summary, our results suggest that in most likely and common forms assortative mating quickly results in the domination of the aging phenotype. The non-aging phenotype prevails only if assortative mating becomes too strong and in case aging and non-aging phenotypes have the same initial fertility. Thus, assortative mating on its own may in fact be the process which partially explains why most species age while some do not.

Our fourth core finding establishes that sexual dimorphisms seem to be more favorable to the aging phenotype than hermaphroditism. In accordance with this result, we are able to make a testable prediction that a lack of senescence should be more common in hermaphroditic organisms. Furthermore, this result is also connected to a finding which suggests that an increase in the cost of the male function in hermaphrodites correlates with an increase of the positive effect of pathogen pressure on the persistence of the aging phenotype. In any case, these results offer novel viewpoints on the evolution of aging and suggest that the common practice of considering haploid hermaphrodites (or even haploid asexuals) as a proxy for diploid true sexuals (i.e. females and males) in population genetics models may not always be appropriate.

Overall, our results suggest that aging may be preferable to non-aging in several situations. Furthermore, these situations are reasonably likely, some of them occur at least in some populations, and others are almost universal (pathogen pressure). Therefore, it seems possible that these scenarios and effects are combined in nature, making aging preferable to non-aging for most species.

## Acknowledgement

The project was supported by the CETOCOEN PLUS (CZ.02.1.01/0.0/0.0/15_003/0000469) project of the Ministry of Education, Youth and Sports of the Czech Republic. The project was also supported by the RECETOX Research Infrastructure (LM2015051 and CZ.02.1.01/0.0/0.0/16_013/0001761). Furthermore, Peter Lenart received support from the Brno Ph.D. Talent. Ludek Berec acknowledges institutional support RVO:60077344.

## Supplementary Information

**Figure S1:**
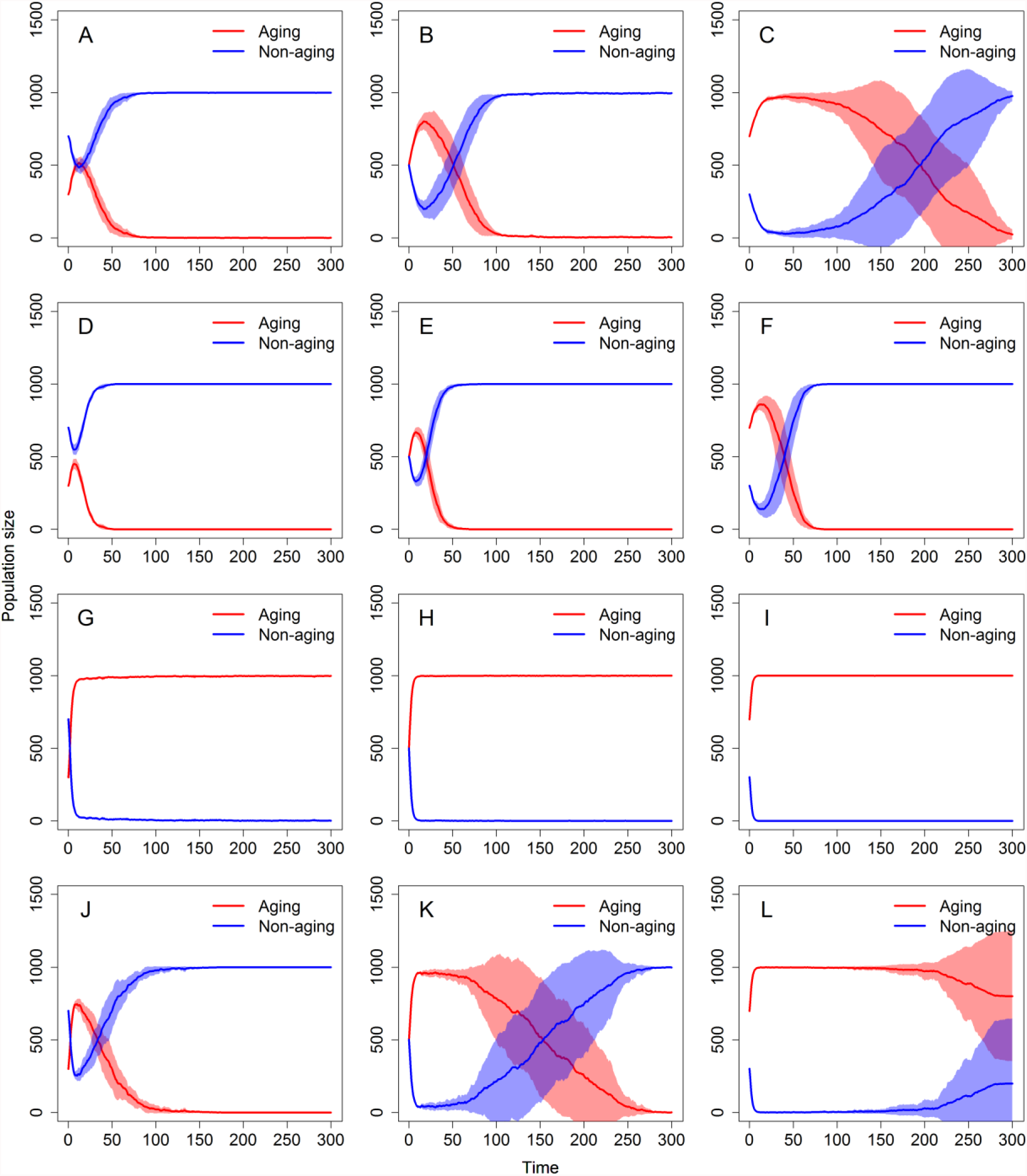
Changes in the representation of aging and non-aging phenotypes in the population under different simulation settings. Panels A) to L) correspond to the respective lines 1 to 12 of Table 2.

**Figure S2:**
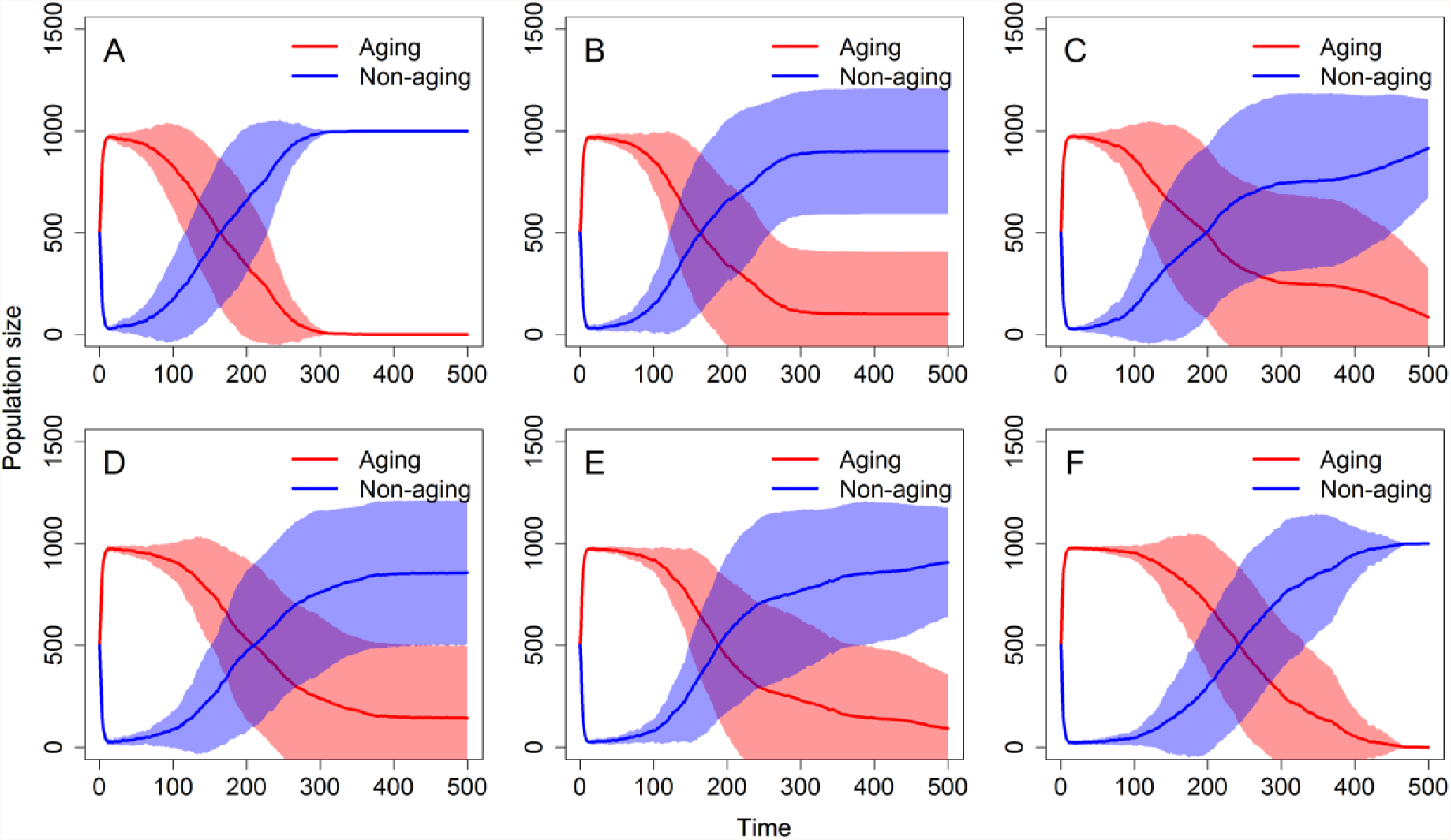
Changes in the representation of aging and non-aging phenotypes in the population under different simulation settings. A) *E_2_* = 0.2, *P =* 1,000. B) *E_2_* = 0.2, *P =* 2,000. C) *E*_*2*_ = 0.2, *P =* 3,000. D) *E*_*2*_ = 0.4, *P =* 1,000. E) *E*_*2*_ = 0.4, *P =* 2,000. F) *E*_*2*_ = 0.4, *P =* 3,000. Other parameters: *n*_*a*_ =4, *n*_*i*_, = 1, *n*_*p*_ = 2.

**Figure S3:**
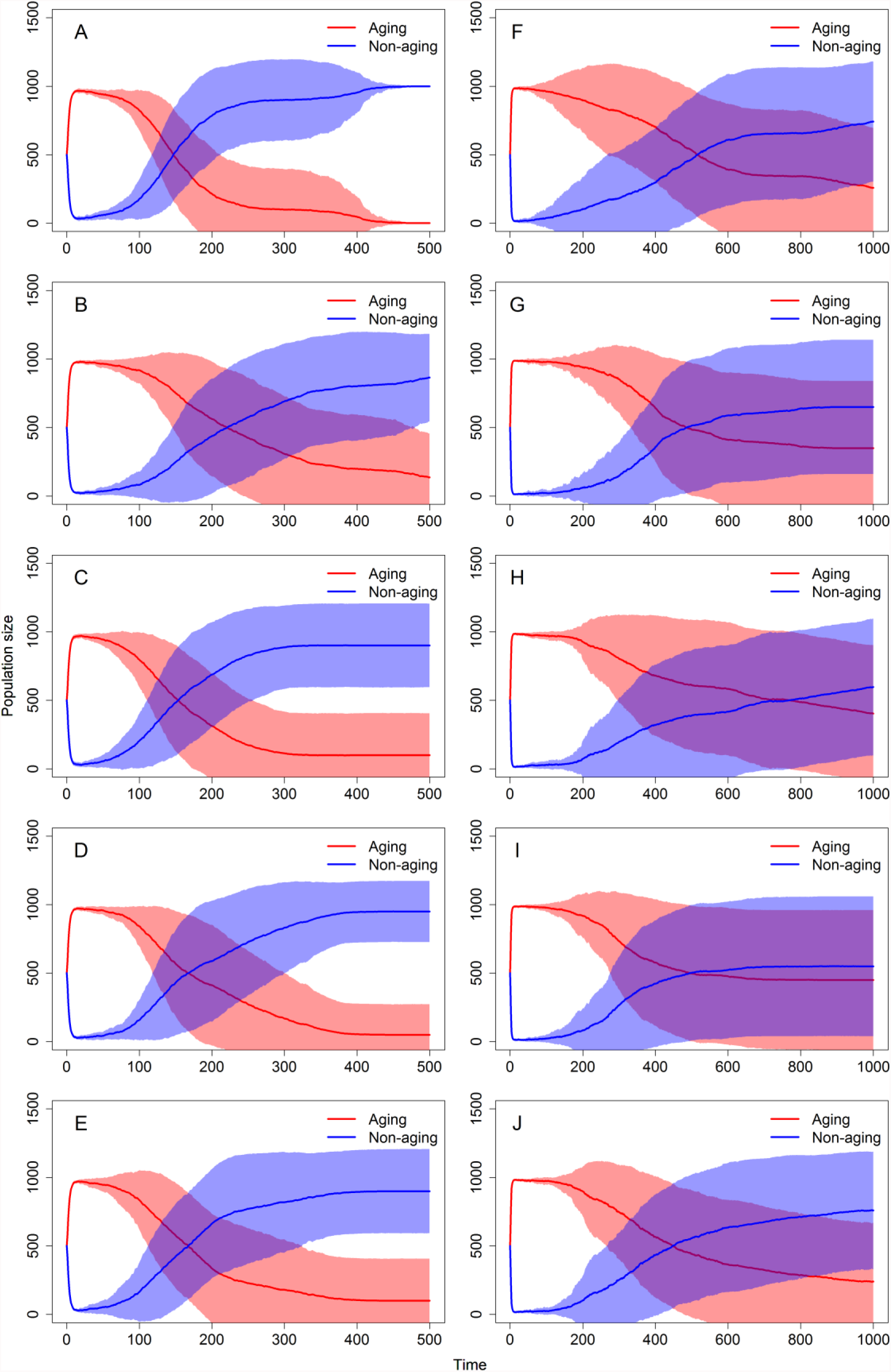
Changes in the representation of aging and non-aging phenotypes in the population with hermaphrodites under different simulation settings. Initial fertility of both aging and non-aging phenotypes is 1.5 for panels A-E and 2 × 1.5 for panels F-J. A and F) No infection; B and G) *n*_*a*_ = 4,*n*_*i*_ = 1; C and H) *n*_*a*_ = 8, *n*_*i*_ = 1; D and I) *n*_*a*_ =4, *n*_*i*_ = 2; E and J) *n*_*a*_ = 8, *n*_*i*_ = 2. Moreover, *E*_*2*_ = 0.4 in all panels with infection.

**Figure S4:**
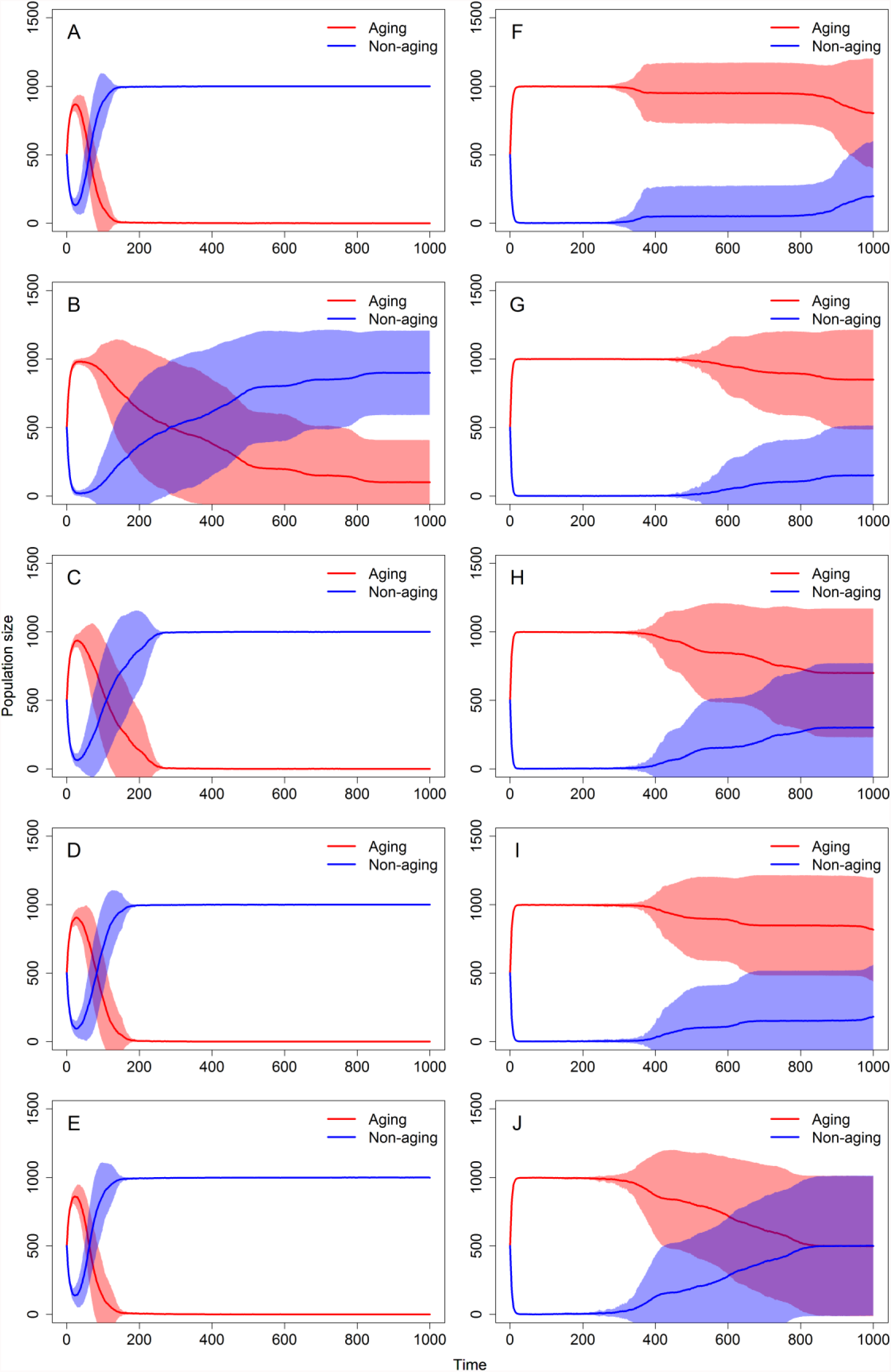
Changes in the representation of aging and non-aging phenotypes in the population with true sexuals under different simulation settings. Initial fertility of both aging and non-aging phenotypes is 1.5 for panels A-E and 2 × 1.5 for panels F-J. A and F) No infection; B and G) *n*_*a*_ = 4, *n*_*i*_ = 1; C and H) *n*_*a*_ = 8, *n*_*i*_ = 1; D and I) *n*_*a*_ =4,*n*_*i*_ = 2; E and J) *n*_*a*_ = 8, *n*_*i*_ = 2. Moreover, *E*_*2*_ = 0.4 in all panels with infection.

## References

1. Kirkwood TBL, Austad SN (2000) Why do we age? Nature 408(6809):233–238.

2. Mart ne E (1998) Mortality Patterns Suggest Lack of Senescence in Hydra. Exp Gerontol 33(3):217–225.

3. Schaible R, et al. (2015) Constant mortality and fertility over age in Hydra. Proc Natl Acad Sci 112(51):15701–15706.

4. Jones OR, et al. (2014) Diversity of ageing across the tree of life. Nature 505(7482):169–173.

5. Jones OR, Vaupel JW (2017) Senescence is not inevitable. Biogerontology 18(6):965–971.

6. Ruby JG, Smith M, Buffenstein R Naked mole-rat mortality rates defy Gompertzian laws by not increasing with age. eLife 7. doi:10.7554/eLife.31157.

7. Weismann A, Poulton EB, Schönland S, Shipley AE (1891) Essays upon heredity and kindred biological problems, by Dr. August Weismann. Ed. by Edward B. Poulton, Selmar Schönland, and Arthur E. Shipley. Authorised translation. (Clarendon Press, Oxford,). 2d ed. Available at: http://www.biodiversitylibrary.org/bibliography/28066.

8. Medawar PB (1952) An Unsolved Problem of Biology: An Inaugural Lecture Delivered at University College, London, 6 December, 1951 (H.K. Lewis and Company).

9. Williams GC (1957) Pleiotropy, Natural Selection, and the Evolution of Senescence. Evolution 11(4):398–411.

10. Kirkwood TB (1977) Evolution of ageing. Nature 270(5635):301–304.

11. Kirkwood TBL, Holliday R (1979) The Evolution of Ageing and Longevity. Proc R Soc Lond B Biol Sci 205(1161):531–546.

12. Kirkwood TBL, Melov S (2011) On the Programmed/Non-Programmed Nature of Ageing within the Life History. Curr Biol 21(18):R701–R707.

13. Longo VD, Mitteldorf J, Skulachev VP (2005) Programmed and altruistic ageing. Nat Rev Genet 6(11):866–872.

14. Mitteldorf JJ (2012) Adaptive aging in the context of evolutionary theory. Biochem Biokhimiia 77(7):716–725.

15. Mitteldorf J, Martins ACR (2014) Programmed Life Span in the Context of Evolvability. Am Nat 184(3):289–302.

16. Skulachev VP (2012) What is “phenoptosis” and how to fight it? Biochem Biokhimiia 77(7):689–706.

17. Libertini G (1988) An adaptive theory of increasing mortality with increasing chronological age in populations in the wild. J Theor Biol 132(2):145–162.

18. Bowles JT (1998) The evolution of aging: a new approach to an old problem of biology. Med Hypotheses 51(3):179–221.

19. Bredesen DE (2004) The non-existent aging program: how does it work? Aging Cell 3(5):255–259.

20. Prinzinger R (2005) Programmed ageing: the theory of maximal metabolic scope. EMBO Rep 6(Suppl 1):S14–S19.

21. Mitteldorf J, Goodnight C (2012) Post-reproductive life span and demographic stability. Oikos 121(9):1370–1378.

22. Mitteldorf J (2006) Chaotic population dynamics and the evolution of ageing. Evol Ecol Res 8(3):561–574.

23. Travis JMJ (2004) The Evolution of Programmed Death in a Spatially Structured Population. J Gerontol A Biol Sci Med Sci 59(4):B301–B305.

24. Mitteldorf J, Pepper J (2009) Senescence as an adaptation to limit the spread of disease. J Theor Biol 260(2):186–195.

25. Martins ACR (2011) Change and Aging Senescence as an Adaptation. PLOS ONE 6(9):e24328.

26. Werfel J, Ingber DE, Bar-Yam Y (2015) Programed Death is Favored by Natural Selection in Spatial Systems. Phys Rev Lett 114(23):238103.

27. Kowald A, Kirkwood TBL (2016) Can aging be programmed? A critical literature review. Aging Cell 15(6):986–998.

28. Lenart P, Bienertová-Vašků J (2017) Keeping up with the Red Queen: the pace of aging as an adaptation. Biogerontology 18(4):693–709.

29. Howard RS, Lively CM (1994) Parasitism, mutation accumulation and the maintenance of sex. Nature 367(6463):554–557.

30. Lively CM, Howard RS (1994) Selection by parasites for clonal diversity and mixed mating. Phil Trans R Soc Lond B 346(1317):271–281.

31. Howard RS, Lively CM (2002) The Ratchet and the Red Queen: the maintenance of sex in parasites. J Evol Biol 15(4):648–656.

32. Howard RS, Lively CM (2003) Opposites attract? Mate choice for parasite evasion and the evolutionary stability of sex. J Evol Biol 16(4):681–689.

33. Agrawal A, Lively CM (2002) Infection genetics: gene-for-gene versus matching-alleles models and all points in between. Evol Ecol Res 4(1):91–107.

34. Dybdahl MF, Jenkins CE, Nuismer SL (2014) Identifying the Molecular Basis of Host-Parasite Coevolution: Merging Models and Mechanisms. Am Nat 184(1):1–13.

35. Borgia G, Blick J (1981) Sexual competition and the evolution of hermaphroditism. J Theor Biol 89(3):523–532.

36. Schumer M, et al. (2017) Assortative mating and persistent reproductive isolation in hybrids. Proc Natl Acad Sci 114(41):10936–10941.

